# Inhibitors of cannabinoid receptor 1 suppress the cellular entry of Lujo virus

**DOI:** 10.1101/2023.05.24.542186

**Authors:** Miyuki Kimura, Risa Matsuoka, Satoshi Taniguchi, Junki Maruyama, Slobodan Paessler, Saori Oka, Atsushi Yamashita, Takasuke Fukuhara, Yoshiharu Matsuura, Hideki Tani

**Author notes:** Corresponding Author: Hideki Tani, Ph.D. Director, Department of Virology, Toyama Institute of Health, 17-1 Nakataikoyama, Imizu-shi, Toyama, 939-0363, Japan. Department of Bacteriology II, National Institute of Infectious Diseases, Tokyo 208-0011, Japan.

## Abstract

Lujo virus (LUJV), which belongs to Mammarenavirus, family *arenaviridae*, has emerged as pathogen causing severe hemorrhage fever with high mortality. Currently, there are no effective treatments for arenaviruses, including LUJV. Here, we screened chemical compound libraries of Food and Drug Administration (FDA)-approved drug and G protein-coupled receptor-associated drugs to identify effective antivirals against LUJV targeting cell entry using a vesicular stomatitis virus-based pseudotyped virus bearing the LUJV envelope glycoprotein (GP). Cannabinoid receptor 1 (CB1) antagonists, such as rimonabant, AM251 and AM281, have been identified as robust inhibitors of LUJV entry. The IC_50_ of rimonabant was 0.26 and 0.53 μM in Vero and Huh7 cells, respectively. Analysis of the cell fusion activity of the LUJV GP in the presence of CB1 inhibitors revealed that these inhibitors suppressed the fusion activity of the LUJV GP. Moreover, rimonabant, AM251 and AM281 reduced the infectivity of authentic LUJV *in vitro,* suggesting that the antiviral activity of CB1 antagonists against LUJV is mediated, at least in part, by inhibition of the viral entry, especially, membrane fusion. These findings suggest promising candidates for developing new therapies against LUJV infections.

**Importance:** To investigate antiviral drugs against Lujo virus (LUJV), we screened chemical compound libraries to identify effective antivirals against LUJV entry. CB1 antagonists were identified as robust inhibitors of LUJV entry. The cell fusion activity of LUJV GP was suppressed by CB1 inhibitors. Furthermore, CB1 antagonists reduced the infectivity of authentic LUJV. These findings suggest promising candidates for developing new therapies against LUJV infections.

## 1. Introduction

Mammarenavirus genus, family *arenaviridae* is classified into Old World (OW) and New World (NW) arenaviruses based on phylogenetic and serological analysis and geographical distribution (Charrel et al., 2008; Wulff et al., 1978). Lassa virus (LASV), an OW arenavirus, causes hemorrhagic fever and is estimated to infect several hundred thousand people in West Africa annually (Centers for Disease Control and Prevention, 2022). NW arenaviruses, such as the Junin (JUNV), Machupo (MACV), Chapare, Sabia, and Guanarito viruses, also cause severe viral hemorrhagic fever with a high fatality (Moraz and Kunz, 2011). Lujo virus (LUJV) was newly identified as a cause of hemorrhagic fever in South Africa, with a fatality rate of 80% (4 cases out of 5) in 2008 (Briese et al., 2009).

Arenaviruses are enveloped viruses that possess a negative-strand RNA that encodes a viral polymerase, nucleoprotein, matrix protein and glycoprotein (GP). Phylogenetic analysis of the amino acid sequences of ORFs revealed that LUJV is uniquely positioned among arenaviruses. In particular, GP sequence analysis revealed that LUJV GP1 relating to interaction with receptor was phylogenetically distant from that of OW and NW arenaviruses (Briese et al., 2009). The entry mechanisms of LUJV is different from those of OW and NW arenaviruses using vesicular stomatitis virus (VSV)-based pseudotyped virus system, which has been developed and has allowed us to investigate the entry mechanisms of highly pathogenic viruses categorized as highly biosafety level (BSL)-3 or BSL-4 pathogens, such as LUJV, in a safe and convenient manner (Tani et al., 2014). OW arenaviruses such as LASV and some of clade C NW arenaviruses employ α-dystroglycan (α-DG) as a cell surface receptor (Cao et al., 1998; Rojek et al., 2007). Many pathogenic viruses in NW arenaviruses use the human transferrin receptor 1 (hTfR1) as the receptor for cell entry (Flanagan et al., 2008; Radoshitzky et al., 2007). LUJV utilizes neither α-DG nor hTfR1 (Tani et al., 2014) but the transmembrane protein neuropilin 2 (NRP2) (Raaben et al., 2017). Arenaviruses exploit the pH-dependent fusion of the viral envelope with the endosomal membrane, mediated by GP, for cellular entry (Di Simone et al., 1994; Tani et al., 2014). Lysosomal-associated membrane protein 1 has been identified as an intracellular receptor for LASV (Jae et al., 2014). CD63, a tetraspanin residing in intracellular vesicles including endosomes and lysosomes, promotes LUJV GP-mediated membrane fusion (Raaben et al., 2017). Recent studies have shown that CD63 is an important factor in determining the host range of LUJV (Saito et al., 2021).

Effective treatments for arenaviruses, including LUJV, are yet to be established. Nucleoside analogs, such as ribavirin and favipiravir, have been demonstrated to be effective against some arenaviruses, such as LASV and JUNV; however, administration of these antiviral agents after symptom onset did not exert their antiviral effects (Gowen et al., 2013; Gowen et al., 2015; Gowen et al., 2007; Mendenhall et al., 2011). Recently, several LUJV inhibitors have been identified by screening chemical compound libraries (Cao et al., 2021; Welch et al., 2021). Here, we screened 1,061 Food and Drug Administration (FDA)-approved drugs and 533 G protein-coupled receptor (GPCR)-associated drugs to identify antiviral reagents targeting the cellular entry of LUJV, using pseudotyped VSV bearing LUJV GP encoding the luciferase gene as a reporter (LUJpv). Cannabinoid receptor 1 (CB1) agonist, such as rimonabant and AM251, were newly identified as robust inhibitors of LUJpv entry. Furthermore, these compounds were confirmed to exhibit potent antiviral activity against authentic LUJV *in vitro*. These results provide novel antivirals against LUJV.

## 2. Materials and Methods

### 2.1. Plasmids and cells

pCAG-LUJV GP, the LUJV GP expressing plasmid was prepared as described previously (Tani et al., 2014). pCAG-T7pol, an expressing plasmid encoding T7 RNA polymerase under the control of the CAG promoter, pT7ECMVLuc, reporter plasmid encoding a firefly luciferase gene under the control of the T7 promoter were used (Tani et al., 2014). The cDNA of CD63 was obtained from CD63-pEGFP C2 plasmid provided from Dr. Paul Luzio (Addgene plasmid #62964) by PCR.

Vero or VeroE6 (African green monkey kidney), Huh7 (human hepatocellular carcinoma), 293T (human embryo kidney) and BHK (baby hamster kidney) cell lines were maintained in Dulbecco’s modified Eagle’s medium (DMEM) (Nacalai tesque, Kyoto, Japan) supplemented with 10% FBS at 37 °C under 5% CO_2_.

### 2.2. Chemicals

The GPCR compound library and FDA-approved drug library were purchased from TargetMol Chemicals Inc. (#L1500 and #L4200, Boston, MA). CB1 inhibitors, i.e., rimonabant, AM251, and AM281, were purchased from Sigma-Aldrich (St. Louis, MI), FUJIFILM Wako Pure Chemical Corporation (Osaka, Japan), and Cayman Chemical (Ann Arbor, MI, USA), respectively. AM630, a cannabinoid receptor type 2 (CB2) inhibitor, was obtained from FUJIFILM Wako Pure Chemical Corporation.

### 2.3. Generation of pseudotyped viruses

Pseudotyped VSVs bearing the LUJV GP or VSV G were prepared as previously described (Tani et al., 2014). Briefly, 293T cells were seeded on collagen I-coated plates and transfected with pCAG-LUJV GP or pCAG-VSV G. At 24 h post-transfection, cells were inoculated with G-complemented VSVΔG/Luc (*G-VSVΔG/Luc) or *G-VSVΔG/GFP at a multiplicity of infection (moi) of 0.5. After 2 h of virus adsorption, the cells were washed four times with serum-free DMEM to remove any unadsorpted virus. At 24 h post-infection, the supernatant containing the pseudotyped virus was harvested and centrifuged to remove cell debris. The resulting supernatant was kept at –80°C until use.

### 2.4. Screening of chemical compound libraries

Vero cells seeded on 96-well tissue culture plate were pre-treated with 5 μM of a compound for 1 h and inoculated with LUJpv or VSVpv. After 24 h, the infectivity of these pseudotyped viruses was determined by measuring luciferase activity using the PicaGene BrillianStar-LT Luciferase Assay System (TOYO B-Net Co., Ltd., Tokyo, Japan) and GloMax Navigator System G2000 (Promega Corporation, Madison, WI, USA) or by observing EGFP expression under a fluorescence microscope (IX71, Olympus, Tokyo, Japan).

### 2.5. Cell viability

Cells were treated with the indicated concentrations of rimonabant, AM251, AM281, or AM630 for 24 h, and cell viability was evaluated by measuring luciferase activity using CellTiter-Glo 2.0 Cell Viability Assay (Promega Corporation) and GloMax Navigator System G2000.

### 2.6. Cell fusion assay

Cell fusion was determined using a previously established quantitative reporter assay (Takikawa et al., 2000; Tani et al., 2014). Briefly, BHK cells were co-transfected with pCAG-LUJV GP and pCAG-T7pol. At 4 h post-transfection, the cells were harvested with trypsin, re-seeded in a 96-well tissue culture plate, and incubated for 24h. The target BHK cells were co-transfected with CD63-pEGFP C2, an expression plasmid encoding CD63 under the control of the CMV promoter, and pT7ECMVLuc, a reporter plasmid encoding a firefly luciferase gene under the control of the T7 promoter. At 24 h post-transfection, the target cells were harvested and co-cultured with cells expressing LUJV GP and T7 RNA polymerase in the presence of the indicated concentrations of rimonabant, AM251, or AM630. After 8 h, co-cultured cells were treated with citrate-phosphate buffer adjusted to pH 4.2 for 2 min, and the buffer was replaced with fresh medium with or without the indicated concentration of rimonabant, AM251, or AM630. After 12 h, the level of cell fusion was determined by measuring the luciferase activity.

### 2.7. Viral yield reduction assay

Vero cells seeded in 12-well tissue culture plates were pre-treated with the indicated concentration of CB1 or CB2 inhibitors for 4 h. The cells were inoculated with authentic LUJV (#200809232Zambia) at an MOI of 0.001. After 30 min, the cells were washed thrice with DMEM containing 2% FBS and incubated with the indicated concentrations of CB1 or CB2 inhibitors. Four days after the infection, the supernatant containing the virus was harvested and were 10-fold serially diluted for viral titration. VeroE6 cells were inoculated with serially diluted virus and incubated for 30 min at 37°C. The viral solution was removed, and the cells were overlaid with Eagle’s MEM supplemented with 1% Avicel RC-591 NF, 100 units/mL penicillin-streptomycin, and 5% FBS. At 7 days post-infection, the cells were fixed with 10% formalin in PBS for 30 min and stained with crystal violet. The viral yield was determined by counting the number of plaques. All experiments on the infectious virus were conducted in a biosafety level 4 (BSL-4) laboratory at the University of Texas Medical Branch (Galveston, TX, USA).

### 2.8 Statistics

Statistical significance was analyzed using an unpaired two-tailed Student’s t-test.

## 3. Results

### 3.1. Screening of chemical inhibitors against LUJV entry

To identify compounds that inhibit LUJV entry, we screened libraries of 1,061 FDA-approved drugs and 533 GPCR-associated compounds using LUJpv. Vero cells were pre-treated with 5 μM of each compound for 1 h, inoculated with LUJpv, and incubated for 24 h. The infectivity of LUJpv were assessed by measuring the luciferase activity. As shown in Table 1, we identified 12 of the 1,594 compounds that suppressed the infectivity of LUJpv to less than 10% of that in control cells treated with 0.5% DMSO. Four of the 12 compounds—rimonabant hydrochloride, AM251, AM281 and rimonabant—were CB1 inhibitors. Therefore, we focused on the inhibitory effects of CB1 antagonists on LUJV entry.

**Table 1.**
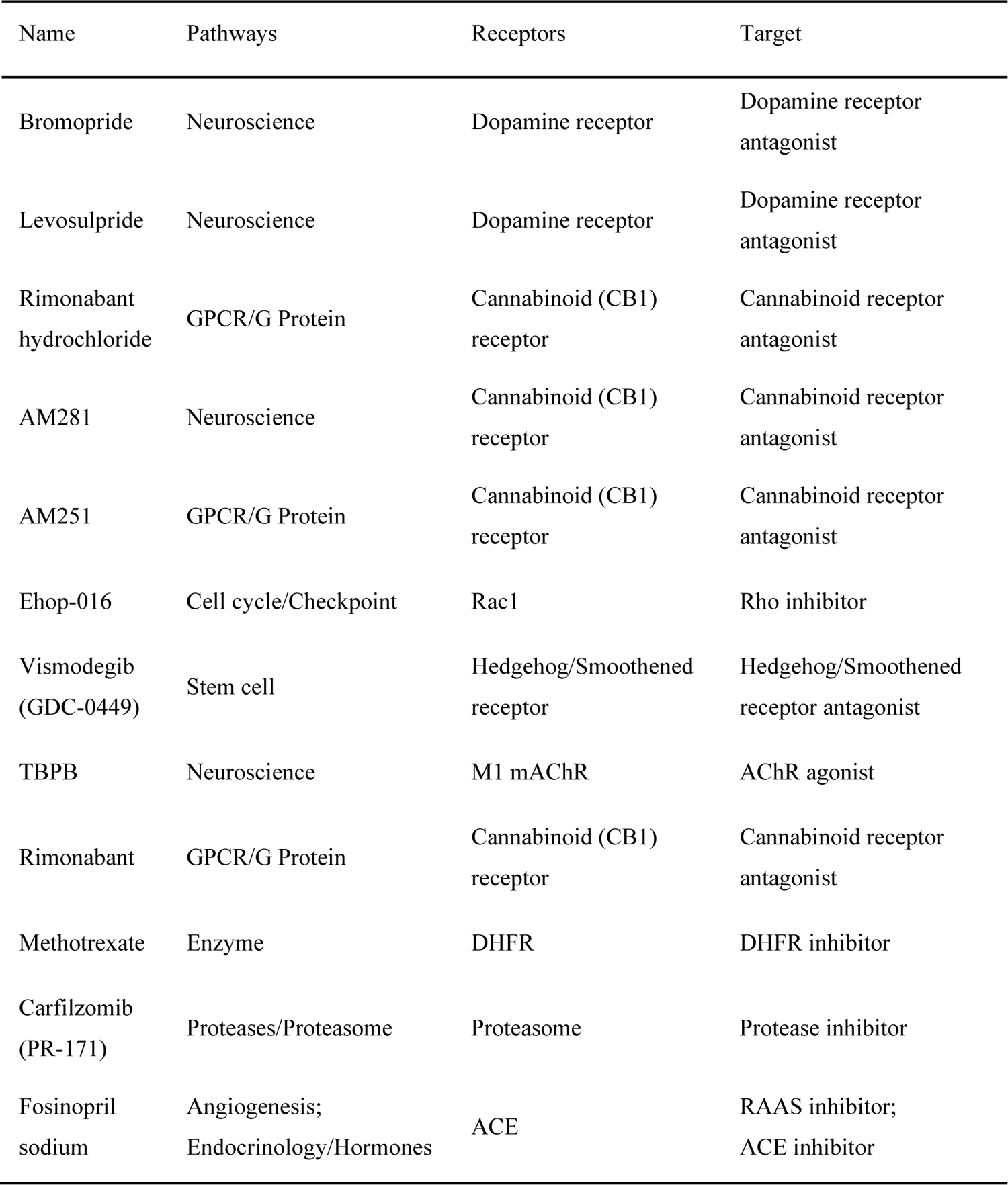

| Name | Pathways | Receptors | Target |
| --- | --- | --- | --- |
| Bromopride | Neuroscience | Dopamine receptor | Dopamine receptor antagonist |
| Levosulpride | Neuroscience | Dopamine receptor | Dopamine receptor antagonist |
| Rimonabant hydrochloride | GPCR/G Protein | Cannabinoid (CB1) receptor | Cannabinoid receptor antagonist |
| AM281 | Neuroscience | Cannabinoid (CB1) receptor | Cannabinoid receptor antagonist |
| AM251 | GPCR/G Protein | Cannabinoid (CB1) receptor | Cannabinoid receptor antagonist |
| Ehop-016 | Cell cycle/Checkpoint | Rac1 | Rho inhibitor |
| Vismodegib (GDC-0449) | Stem cell | Hedgehog/Smoothened receptor | Hedgehog/Smoothened receptor antagonist |
| TBPB | Neuroscience | M1 mAChR | AChR agonist |
| Rimonabant | GPCR/G Protein | Cannabinoid (CB1) receptor | Cannabinoid receptor antagonist |
| Methotrexate | Enzyme | DHFR | DHFR inhibitor |
| Carfilzomib (PR-171) | Proteases/Proteasome | Proteasome | Protease inhibitor |
| Fosinopril sodium | Angiogenesis;<br>Endocrinology/Hormones | ACE | RAAS inhibitor;<br>ACE inhibitor |

### 3.2. Inhibitory effect of CB1 inhibitors on the infectivity of LUJpv

The two types of cannabinoid receptors, CB1 and CB2, are seven-transmembrane G protein-coupled receptors that regulate a wide range of physiological functions (Cabral et al., 2015; Galiègue et al., 1995; Katona and Freund, 2012; Pertwee, 2015). To confirm the inhibitory effect of CB1 inhibitors on the infectivity of LUJpv, Vero cells were pre-treated with the CB1 inhibitors (rimonabant, AM251 and AM281) or the CB2 inhibitor AM630 at the indicated concentration for 1 h and inoculated with LUJpv or VSVpv. At 24 h post-infection, infectivity was examined by measuring luciferase activity. The infectivity of LUJpv were inhibited by all three CB1 inhibitors in a dose-dependent manner and were reduced to approximately 2% at 4 μM of rimonabant, 3% at 5 μM of AM251, and 5% at 5 μM of AM281, in contrast, that of VSVpv were not (Fig. 1, graphs). We also examined the effect of CB1 inhibitors on the infectivity of LUJpv, which encodes GFP as a reporter. The number of cells expressing GFP was suppressed by the CB1 inhibitors in a dose-dependent manner, which correlated with luciferase activity (Fig. 1, lower panels). A decrease in LUJpv infection induced by the CB1 inhibitors was also observed in Huh7 and 293T cells (data not shown). The value of IC_50_ of rimonabant and AM251 for Vero cells were lower than those for Huh7 cells, and rimonabant was more effective in both cell lines, with selective index (SI) values >78 in Vero cells and >38 in Huh7 cells (Table 2).

**Fig. 1.**
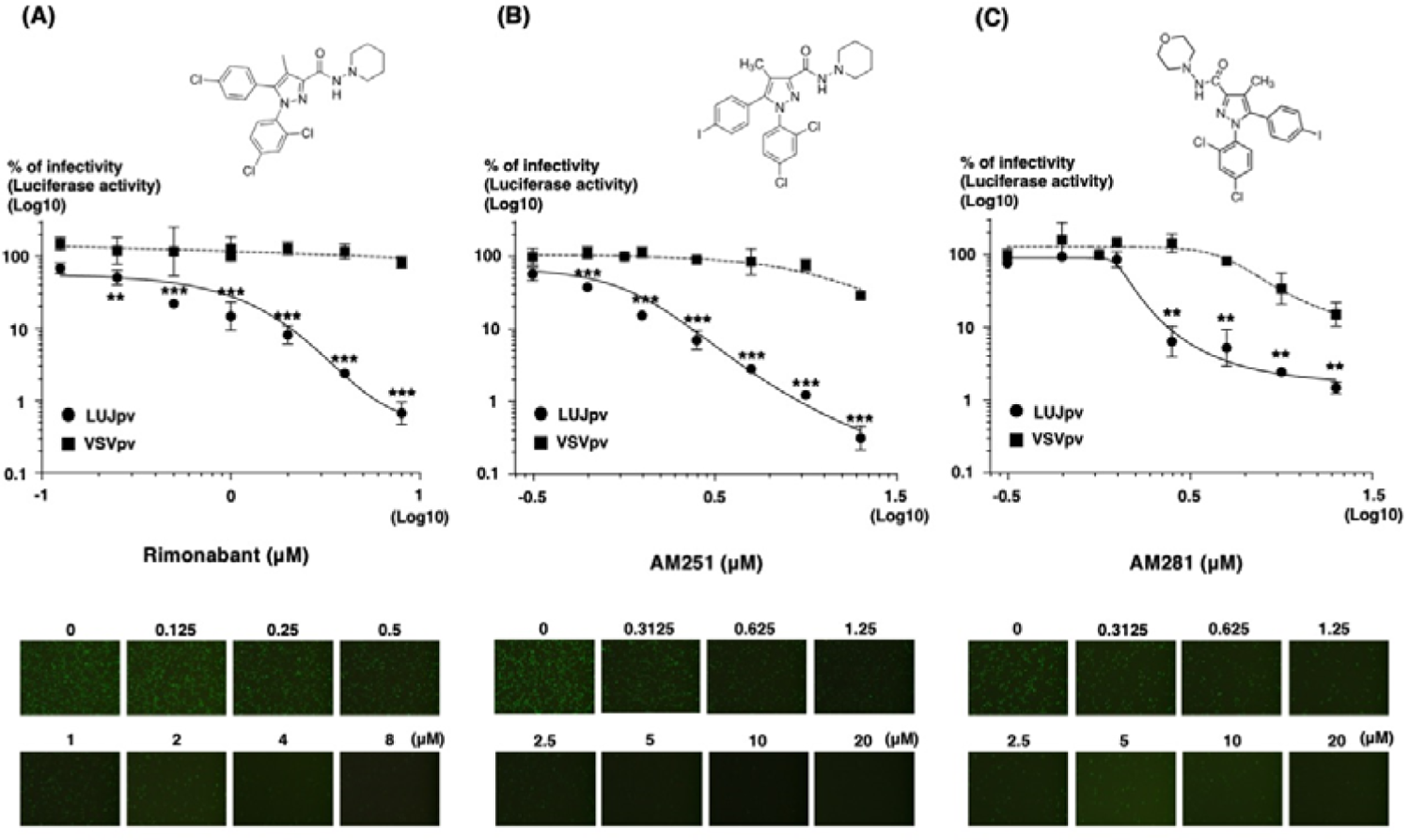
Inhibition of LUJpv infection by CB1 antagonists. Vero cells were pre-treated with the indicated concentrations of rimonabant (A), AM251 (B), or AM281 (C) for 1 h and inoculated with LUJpv encoding luciferase, GFP as a reporter, or VSVpv encoding luciferase. Infectivity was determined by measuring luciferase activity or by fluorescence microscopic observation of GFP expression at 24 h post-infection. The structural formula of rimonabant, AM251, and AM281 are described in panel A, B, and C, respectively. Black circles and squares indicate the infectivity of LUJpv and VSVpv, respectively. Results are from three independent assays, with error bars representing the standard deviations. The statistical significance of the differences between the infectivity of pseudotyped viruses with and without the inhibitor was analyzed using Student’s t-test (** p<0.005, ***p<0.001).

**Table 2.**

| Cells | Inhibitor | IC<sub>50</sub> (μM) | CC<sub>50</sub> (μM) | SI |
| --- | --- | --- | --- | --- |
| Vero | rimonabant | 0.26 | >20 | >78 |
|  | AM251 | 0.41 | 19.94 | 41 |
| Huh7 | rimonabant | 0.53 | >20 | >38 |
|  | AM251 | 1.71 | >20 | >12 |

In contrast, statistical analysis showed that the infectivity of LUJpv was significantly suppressed by the CB2 inhibitor AM630. However, the inhibitory effect of the CB1 inhibitors was remarkably higher than that of the CB2 inhibitor (Fig. 2). The infectivity of VSVpv was unaffected by the CB2 inhibitor (Fig. 2). Cytotoxicity of CB1 inhibitors was not observed except for at 20 μM of AM251 in Vero cells (Fig. 3). These results suggest that the CB1 inhibitors, rimonabant, AM251 and AM281 reduce cell entry of LUJV.

**Fig. 2.**
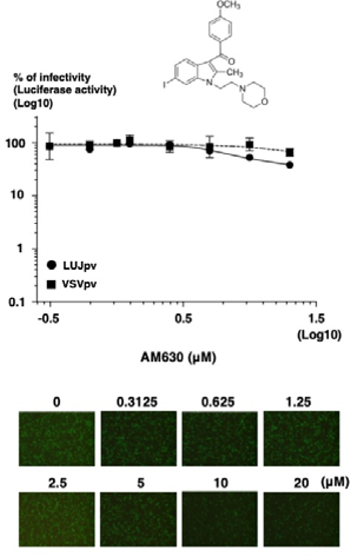
Effect of CB2 inhibitor on the infectivity of LUJpv Vero cells were pre-treated with the indicated concentrations of AM630 for 1 h and inoculated with LUJpv encoding luciferase, GFP gene as a reporter, or VSVpv encoding the luciferase gene. Infectivity was determined by measuring luciferase activity or by fluorescence microscopic observation of GFP expression at 24 h post-infection. The structural formula of AM630 is given above the graph. Black circles and squares indicate the infectivity of LUJpv and VSVpv, respectively. Results are from three independent assays, with error bars representing the standard deviations. The statistical significance of the differences between the infectivity of pseudotyped viruses with and without the inhibitor was analyzed using Student’s t-test (** p<0.005).

**Fig. 3.**
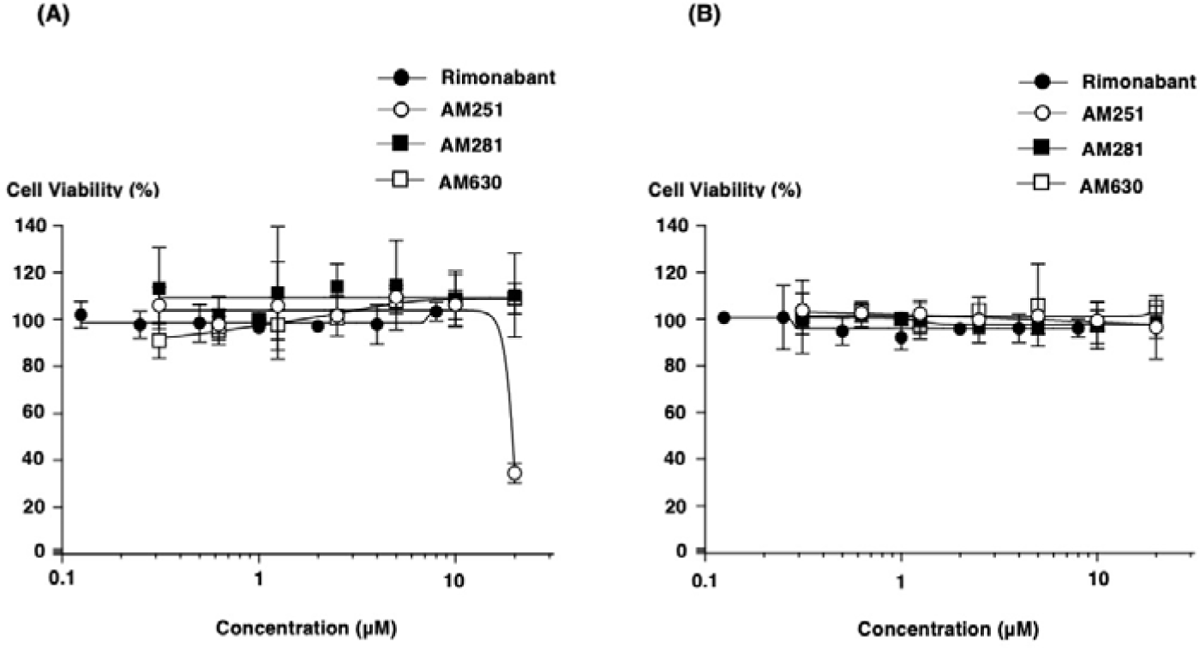
Viability of cells treated with CB1 and CB2 inhibitors Vero cells (A) or Huh7 cells (B) were incubated with the indicated concentrations of rimonabant, AM251, AM281, or AM630 for 24 h. Next, the CellTiter-Glo 2.0 Cell Viability Assay kit was added to the cells and cell viability was determined by measuring luciferase activity. Luciferase activity of the untreated cells was set as 100%. Closed circles, open circles, closed squares, and open squares represent cell viability after treatment with rimonabant, AM251, AM281 and AM630, respectively. Results are from three independent assays, with error bars representing the standard deviations.

### 3.3. Effect of CB1 inhibitors on cell fusion activity of LUJV GP via CD63

To identify the mechanisms by which CB1 antagonists block the entry of LUJV, the effect of CB1 inhibitors on the fusion activity of LUJV GP via CD63, an endosomal receptor of LUJV, was assessed using a quantitative cell fusion assay based on reporter gene activation. A schematic representation of the fusion assay is shown in Figure 4A. Cell fusion was induced in LUJV GP-expressing cells by treatment with buffer adjusted to pH 4.2 in the presence of CD63. CB1 antagonists, rimonabant and AM251 inhibited the cell fusion in a dose-dependent manner, and the fusion activity was reduced to approximately 40% at 5 μM. In contrast, the CB2 inhibitor AM630 did not (Fig. 4B). CB1 antagonists inhibited fusion activity; however, their efficacy was lower than that of LUJpv. Thus, CB1 inhibitors may suppress other step(s) in LUJV entry in addition to membrane fusion.

**Fig. 4.**
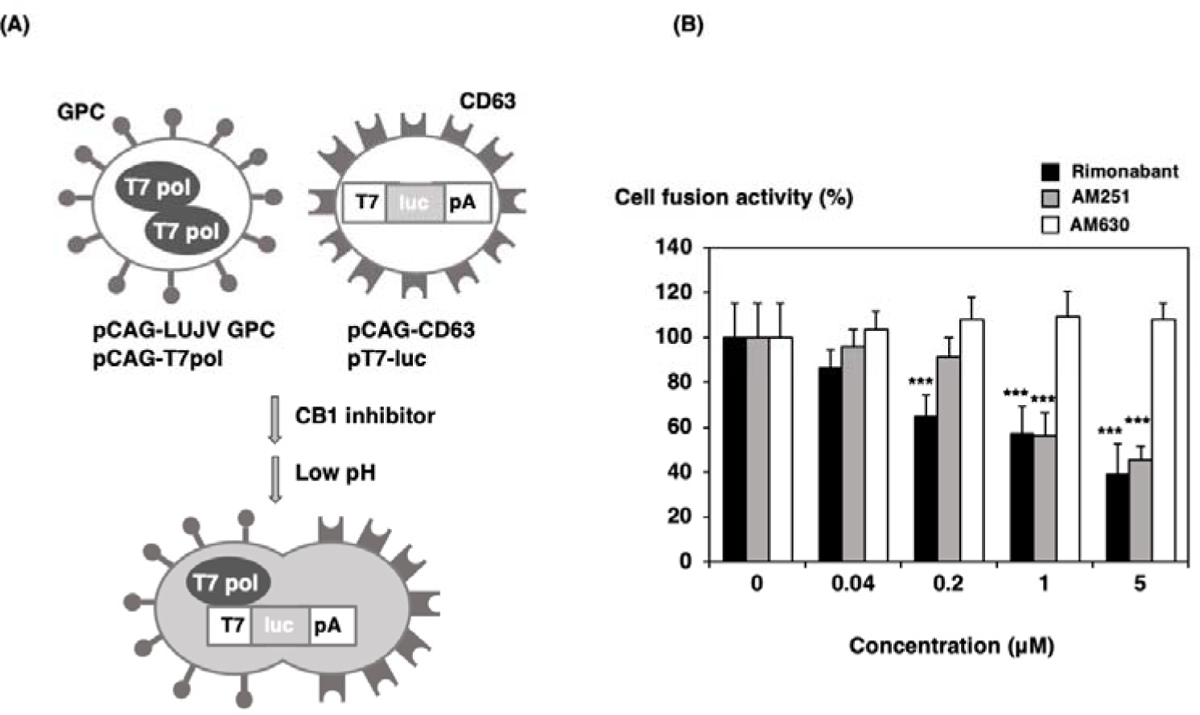
Effect of CB1 inhibitors on the fusion activity of LUJV GP (A) Schematic diagram of the cell fusion assay. (B) Effect of CB1 inhibitors on the fusion activity. BHK cells transfected with pCAG-LUJV GP and pCAG-T7 polymerase were co-cultured with the cells transfected with pCAG-CD63 and luciferase gene under the control of T7 promotor in the presence of the indicated concentration of rimonabant, AM251 or AM630 for 8 h. The cells were treated with the buffer adjusted to pH 4.2 for 2 min, following which the buffer was replaced with fresh medium containing the inhibitor. The cells were incubated for 12 h and the fusion activity was determined by measuring luciferase activity. The fusion activity without inhibitor was set as 100%. Black, gray, and white bars represent the fusion activity in the presence of rimonabant, AM251 and AM630, respectively. The results shown are from three independent assays, with error bars representing standard deviations. Statistical significance of the differences between the infectivity of pseudotyped viruses with and without the inhibitor was analyzed by Student’s t-test (*** p<0.001).

### 3.4. Anti-LUJV activity of CB1 antagonists

Finally, the antiviral activity of CB1 antagonists against authentic LUJV *in vitro* was evaluated. Vero cells pre-treated with CB1 or CB2 inhibitors were inoculated with LUJV in the presence of CB1 or CB2 inhibitors. At 4 days post-infection, the supernatant was harvested, and viral titers were examined. The viral yield in Vero cells treated with the CB1 inhibitors, rimonabant, AM251 and AM281 decreased in a dose-dependent manner. In contrast, the CB2 inhibitor AM630 did not affect the viral yield (Fig. 5). These results suggest that the CB1 antagonists, rimonabant, AM251 and AM281 inhibited the infection of authentic LUJV.

**Fig. 5.**
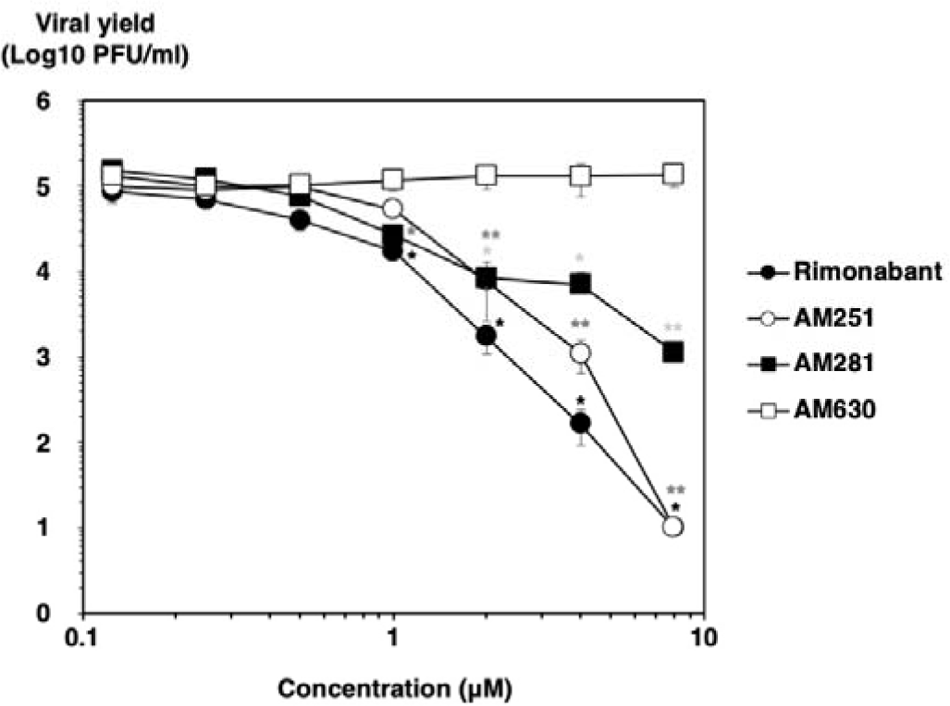
Antiviral activity of CB1 inhibitors against authentic LUJV Vero cells pre-treated with the indicated concentrations of CB1 or CB2 inhibitors for 4 h were inoculated with LUJV at an MOI of 0.001, and incubated at 37 °C in the presence of the indicated concentrations of CB1 or CB2 inhibitors. Four days after the infection, the supernatant was subjected to viral titration. VeroE6 cells were inoculated with serially diluted supernatants and incubated at 37 °C. Five days post-infection, the viral yield was determined by counting the number of plaques. Statistical significance of the differences between the infectivity of pseudotyped viruses with and without the inhibitor was analyzed using Student’s t-test (*p<0.05, **p<0.005, *** p<0.001). Asterisks in black, dark gray, and light gray indicate the results of Student’s t-test for the viral titer with rimonabant, AM251 and AM281, respectively.

## 4. Discussion

The antiviral effects of CB1 inhibitors have been reported previously. Production of hepatitis C virus (HCV) was inhibited by CB1 antagonists (Shahidi et al., 2014). Suppression of hepatitis B virus (HBV) RNA transcription by CB1 inhibitors via inhibiting hepatocyte nuclear factor 4α which is known to stimulate viral RNA synthesis has been described (Sato et al., 2020). CB1expression is induced in patients with chronic HCV infection (Van der Poorten et al., 2010). In this study, we found that LUJV entry was inhibited by CB1 antagonists, which may be due to mechanisms different from those of CB1 antagonists in the replication of HBV and HCV.

Arenavirus fusion inhibitors have been previously described. Some small chemical molecules have been shown to block GP-mediated membrane fusion of arenavirus by binding at the stable signal peptide-GP2 interface, which is the region of GP that triggers membrane fusion. The binding properties of compounds determine their selectivity for OW and NW arenaviruses (Gowen et al., 2021; Plewe et al., 2019; Shankar et al., 2016; York et al., 2008). TRAM-34, a clotrimazole derivative, suppresses the entry of arenaviruses, including LUJV, but not LASV. In MACV and lymphocytic choriomeningitis virus, it has been shown to inhibit the GP fusion (Torriani et al., 2019). Recently, LUJV inhibitors have been screened. Brequiner, selective inhibitor of dihydroorotate dehydrogenase which is believed to act by blocking de novo pyrimidine biosynthesis, 2’-deoxy-2’-fluorocytidine, nucleotide analog and AVN-944, inosine-5’-monophosphate dehydrogenase inhibitors, which are predicted to target viral replication have been described as effective inhibitors for LUJV with SI value of more than 50 (Welch et al., 2021). Trametinib, a MAPK inhibitor, suppresses the entry of LUJV and other pathogenic arenaviruses (Cao et al., 2021). Trametinib targets the GP2 transmembrane domain to inhibit membrane fusion. Our preliminary data showed that rimonabant and AM251 suppressed the infectivity of LUJpv in 293T cells, in which CB1 was not expressed. In addition, 2-arachidonoyl-glycerol, a CB1 agonist, did not enhance LUJpv in Vero cells. These findings implied that CB1 does not necessarily participate in LUJpv infection. Although further studies are needed to identify the mechanisms by which CB1 antagonists inhibit cell fusion, the binding of CB1 inhibitors to the GP region during membrane fusion may be a possible mechanism. Analyses of the effect of CB1 inhibitors on the LUJV fusion using GP mutants may help clarify this possibility in future studies. CB1 antagonists inhibited cell fusion; however, the inhibitory effect was lower than that on the infectivity of LUJpv, suggesting that CB1 antagonists suppress other step(s) in the entry of LUJV, in addition to cell fusion.

In conclusion, we screened chemical compound libraries and identified CB1 antagonists as robust inhibitors of LUJV entry. These antagonists repressed the cell fusion activity of LUJV. Furthermore, the antiviral effects of these inhibitors on the infectivity of authentic LUJV were observed *in vitro*. Although the mechanisms by which CB1 antagonists block the entry of LUJV remain unclear, the inhibitory effect on LUJV entry seems to be mediated, at least in part, by the inhibition of the fusion activity of LUJV GP. Rimonabant induces weight loss and reduces dyslipidemia in obese rodents (Gary-Bobo et al., 2007; Ravinet Trillou et al., 2003) and humans (Després, 2007; Pi-Sunyer et al., 2006; Van Gaal et al., 2005). Rimonabant was approved as a therapeutic drug for obesity in Europe in 2006; however, marketing authorization was suspended in 2008 due to adverse psychiatric side effects. Although CB1 antagonists, such as rimonabant, have been shown to be potent inhibitors of LUJV entry, some improvement, likely to avoid blood-brain barrier penetration, will be needed for the use of these inhibitors as therapeutic agents without side effects.

## Acknowledgements

We are grateful to Mr. Yoshihiro Yoshida and Ms. Kaoru Hounoki for their technical and secretarial assistance.

## Funding

This work was supported by the Japan Agency for Medical Research and Development (AMED) (Grant No. JP19fm0208101) (Y. M.), the National Institutes of Health (Grant No. U01AI151801) (S. P.), and JSPS KAKENHI Grant Number JP15K08510 (H. T.).

## Author contributions

Conceptualization and methodology: M. K., S. T., A. Y., T. F., and H. T.; investigation: M. K., R. M., S. T., J. M., and H. T.; writing–original draft presentation: M. K., S. T., and H. T.; writing–review and editing: M. K., R. M., S. T., S. P., A. Y., T. F., Y. M., and H. T.; supervision, S. P., Y. M., and H. T.; funding acquisition, S. P., Y. M., and H. T. All authors have read and agreed to the published version of the manuscript.

